# Sanfetrinem, an oral β-lactam antibiotic repurposed for the treatment of tuberculosis

**DOI:** 10.1101/2024.10.10.617558

**Authors:** Santiago Ramón-García, Rubén González del Río, María Pilar Arenaz-Callao, Helena Boshoff, Joaquín Rullás, Sara Anca, Mónica Cacho Izquierdo, Esther Porras de Francisco, Esther Pérez Herrán, Angel Santos-Villarejo, Alfonso Mendoza-Losana, Santiago Ferrer-Bazaga, Charles J. Thompson, David Barros Aguirre, Robert H. Bates

**Affiliations:** Department of Microbiology, Radiology, Pediatrics and Public Health, Faculty of Medicine, University of Zaragoza, Zaragoza, Spain; Research & Development Agency of Aragón (ARAID) Foundation, Zaragoza, Spain; Department of Microbiology & Immunology, Life Sciences Centre, University of British Columbia, Vancouver, B.C., Canada; Global Health Medicines R&D, GSK, Tres Cantos, Madrid, Spain; Tuberculosis Research Section, NIH, Bethesda, USA; Discovery DMPK, GSK, Tres Cantos, Madrid, Spain

**Author notes:** Certest Biotec, San Mateo de Gállego, Zaragoza, Spain (Esther Pérez Herrán). Dep. Bioengineering, University Carlos III of Madrid, Spain.

**Keywords:** sanfetrinem, repurposing, tuberculosis, β-lactams, intracellular

## Abstract

Tuberculosis (TB) is historically the world’s deadliest infectious disease. New TB drugs that can avoid pre-existing resistance are desperately needed. The β-lactams are the oldest and most widely used class of antibiotics to treat bacterial infections but, for a variety of reasons, they were largely ignored until recently as a potential treatment option for TB. Recently, a growing body of evidence indicates that later-generation carbapenems in the presence of β-lactamase inhibitors could play a role in TB treatment. However, most of these drugs can only be administered intravenously in the clinic.

We performed a screening of β-lactams against intracellular *Mycobacterium tuberculosis (Mtb)* and identified sanfetrinem cilexetil as a promising oral β-lactam candidate. Preclinical *in vitro* and *in vivo* studies demonstrated that: (i) media composition impacts the activity of sanfetrinem against *Mtb*, being more potent in the presence of physiologically relevant cholesterol as the only carbon source, compared to the standard broth media; (ii) sanfetrinem shows broad spectrum activity against *Mtb* clinical isolates, including MDR/XDR strains; (iii) sanfetrinem is rapidly bactericidal *in vitro* against *Mtb* despite being poorly stable in the assay media; (iv) there are strong *in vitro* synergistic interactions with amoxicillin, ethambutol, rifampicin and rifapentine and, (v) sanfetrinem cilexetil is active in an *in vivo* model of infection. These data, together with robust pre-clinical and clinical studies of broad-spectrum carbapenem antibiotics carried out in the 1990s by GSK, identified sanfetrinem as having potential for treating TB and catalyzed a repurposing proof-of-concept Phase 2a clinical study (NCT05388448) currently underway in South Africa.

## INTRODUCTION

Tuberculosis (TB) is historically the world’s deadliest infectious disease and the COVID-19 pandemic has caused a reversal in the gains made in diagnosing and treating TB globally [1]. Adding to these challenges, drug resistant forms of TB continue to spread, including strains resistant to bedaquiline, a drug approved just over a decade ago that is now the backbone for most MDR treatment regimens [2]. Therefore, new TB drugs that can avoid pre-existing resistance are desperately needed to combat this deadly disease.

TB is caused by a bacterium, *Mycobacterium tuberculosis (Mtb)*. Thus, it is at first surprising to realize that the oldest and most widely used class of antibiotics for bacterial infections, the β-lactams, was largely ignored for decades as a potential treatment option for TB [3]. A multitude of reasons for this historical trend exist, including: (i) *Mtb* constitutively expresses a β-lactamase, BlaC, that efficiently hydrolyzes most β-lactam drugs rendering them inactive [4–6]; (ii) early clinical trials that showed only modest results even in the presence of clavulanic acid [6] and (iii) the fact that the most efficacious β-lactam drugs showing antitubercular activity are not orally bioavailable making them non-optimal for treating TB [7].

However, since the early 2000’s, there has been a growing body of evidence indicating that later-generation carbapenems show potent activity against *Mtb*, particularly in the presence of a β-lactamase inhibitor [8–11]. These results helped inspire a Phase 2a Early Bactericidal Activity (EBA) clinical trial of meropenem with Augmentin (amoxicillin/clavulanate) that demonstrated a robust reduction in sputum CFU counts, similar to the standard therapy (RHZE), definitively proving that β-lactams are active against *Mtb* in human lungs [12]. Unfortunately, meropenem suffers from two of the major drawbacks mentioned above: it requires co-dosing with a β-lactamase inhibitor and can only be administered by intravenous injection. With these limitations, the clinical use of meropenem for TB is largely limited to rescue therapy for hospitalized M/XDR patients [13]. Therefore, an alternative carbapenem drug that could overcome these issues would represent a significant advance.

To this end, and in parallel to other approaches [3], we set out to screen β-lactams for activity in an intramacrophage *Mtb* assay, hypothesizing that compounds able to penetrate and kill mycobacteria within macrophages would have an inherent advantage for TB. This screen identified sanfetrinem and its ester prodrug, sanfetrinem cilexetil, as promising candidates. These compounds were discovered and developed by Glaxo Wellcome in the 1990s as potential broad-spectrum antibiotics [14–16]. Development was terminated following successful Phase 2 trials primarily due to commercial considerations, meaning that sanfetrinem cilexetil could offer an attractive re-purposing opportunity of a clinically validated asset. Here, we present the identification and pre-clinical characterization of this drug against M. tuberculosis.

## MATERIAL AND METHODS

Bacterial strains, general growth conditions and reagents. Mycobacterium strains were routinely propagated at 37°C in Middlebrook 7H9 broth (Difco) supplemented with 10% Middlebrook albumin-dextrose-catalase (ADC)(Difco), 0.2% glycerol and 0.05% (vol/vol) tyloxapol (complete 7H9) or on Middlebrook 7H10 agar plates (Difco) supplemented with 10% (vol/vol) oleic acid-albumin-dextrose-catalase (OADC) (Difco). Hygromycin B (Sigma) was added to the medium (50 µg/mL) to ensure plasmid maintenance when propagating the M. tuberculosis H37Rv-luc, M. tuberculosis H37Rv-gfp and M. bovis BCG-gfp strains. M. tuberculosis H37Rv-luc constitutively expressed the luciferase luc gene from Photinus pyralis (GenBank Accession Number M15077) cloned in a mycobacterial shuttle plasmid derived from pACE-1 [17], while M. tuberculosis H37Rv-gfp and M. bovis BCG-gfp strains harbored the pFPV27 plasmid [18]. M. tuberculosis clinical isolates were provided by the Vall d’Hebron Hospital, Barcelona, Spain and the Tuberculosis Research Section, NIH, Bethesda, USA. Reagents used, including sanfetrinem, other β-lactams and anti-TB drugs, were provided by GSK. Potassium clavulanate was obtained from Fluka (Ref. 33454). Due to compound instability, β-lactams were freshly dissolved from solid stocks on the same day of the experiment.

Extracellular susceptibility assays. Stock solutions of compounds were prepared fresh on the same day of plate inoculation, dissolved in DMSO and dispensed using a HP D3000 Digital Dispenser and HP T8 Dispenserhead Cassettes (Ref No. CV081A) in two-fold serial dilutions in a 384-well plate format. Dose-response assays were performed essentially as previously described [19]. Briefly, 7H9-based broth medium was supplemented with 0.2% glycerol and 10% ADC without detergent (i.e., tween 80 or tyloxapol). Glucose medium was prepared from 7H9 salts with a final concentration of 55 mM glucose. When needed, bovine serum albumin (BSA) was added at 0.5% from a 5 mg/mL solution. Mycobacterial cells were grown to an OD_600_= 0.5-0.8 and stocks were frozen at −80°C. Upon thawing, cells were diluted in assay medium to a final concentration of 10^5^ cells/mL (or 10^6^ cells/mL for the cholesterol assays) and 50 µL/well dispensed. Plates were placed in sealed boxes to prevent from drying of the perimetric wells and incubated without shaking at 37°C for 6 days before addition of a mix solution of 20% tween 80 plus MTT [3-(4,5-dimethylthiazol-2-yl)-2,5-diphenyl tetrazolium bromide] (Stock 5 mg/mL, Acros Organics, Ref. 15224654) or the Bactiter-Glo Luciferase Assay System (Promega, Madison, WI), both used as cell growth indicators. For the MTT read-out, the optical density at 580 nm (OD_580_) was measured in a Spectramax M5 (Molecular Devices) reader using black 384-microclear plate (Ref. 781091, Greiner). For the Bactiter-Glo system, ATP production was measured by luminescence (according to manufacturer indications) in an Envision Multilabel Plate Reader (PerkinElmer) using a white opaque 384-plate (Ref. 781075, Greiner) with Ultra-Sensitive luminescence mode and a measurement time of 50 ms/well. Dose response curves were plotted as percentage of growth compared to untreated internal controls (i.e., wells with no drug added/DMSO control). Moxifloxacin was used as a dose response compound control with 2-fold dilutions starting at 1 μg/mL. For the cholesterol assays, cholesterol was brought into solution (100 mM) by frequent vortexing and heating at 65°C in ethanol-tyloxapol (1:1 v/v). A 1/1,000 dilution was then added to 7H9-based broth medium to give a final concentration of 0.1 mM cholesterol. Then, susceptibility assays on cholesterol as the sole carbon source were performed as previously described [19, 20].

Intracellular susceptibility assays. Up to three independent readouts were used to study the intracellular activity of test compounds: luminescence, fluorescence confocal microscopy and CFU enumeration.

i. Luminescence. The assay was performed essentially as previously described [17]. Frozen stocks of macrophage THP1 cells (ATCC TIB-202) were thawed in RPMI-1640 medium (Sigma) supplemented with 10% fetal bovine serum (Gibco), 2 mM L-glutamine (Sigma) and 1 mM sodium pyruvate (Sigma). THP1 cells were passaged only 5 times without antibiotics and maintained between 2-10×10^5^ cells/mL at 37°C in a humidified, 5% CO_2_ atmosphere. THP1 cells (3×10^8^) were simultaneously differentiated with phorbol-myristate acetate (PMA, 40 ng/mL, Sigma) and infected for 4 hours at a multiplicity of infection (MOI) of 1:1 with a single cell suspension of M. tuberculosis H37Rv-luc cells. After incubation, infected cells were washed four times to remove extracellular bacilli and resuspended in fresh RPMI medium. Infected cells were finally resuspended (2×10^5^ cells/mL) in RPMI medium supplemented with 10% fetal bovine serum (Hyclone), 2 mM L-glutamine and pyruvate and dispensed in white, flat bottom 384-well plates (Greiner) at a concentration of ca. 10,000 cells per well in a final volume of 50 µL (max. 0.5% DMSO). Plates were incubated for 5 days at 37°C under 5% CO_2_ atmosphere, 80% relative humidity, before growth assessment using the Bright-Glo Luciferase Assay System (Promega, Madison, WI) as above described. Internal wells containing drug-free medium with and without infected macrophages established maximum and minimal luminescence production, respectively. A 90% reduction in light production was considered growth inhibition (IC_90_). Every drug or drug combination was assayed in at least three independent experiments.
ii. CFU enumeration. THP1 cells were treated and infected with M. tuberculosis H37Rv cells as above described at an MOI of 1:3 and seeded (50,000 cells/well) onto 24-well plates. Dose range concentrations of sanfetrinem were added at days 1 and 4 post-infection. Macrophages were lysed at days 1, 4 and 6 by osmotic pressure in a V_F_= 500 µL. Then, 100 µL were 10-fold serially diluted in 1x PBS buffer with 0.1% tyloxapol and plated in 7H10 plates. Agar plates were incubated at 37°C for 14-21 days and CFUs enumerated. Plates were checked again at 4 weeks of incubation to count late growers. Cell density was reported as log_10_CFU/mL. The human biological samples were sourced ethically, and their research use was in accord with the terms of the informed consents under an IRB/EC approved protocol.

Drug interaction assays. Checkerboard extracellular assays were used to identify pairwise interaction profiles of sanfetrinem and meropenem against a panel of clinically approved antibiotics and other antimicrobials with known mode of action. Drug activity was determined in 384-well plate format using the MTT or ATP assay, as described above. The fractional inhibitory concentration (FIC) for each compound was calculated as previously described [21]. FICI values indicated the degree of interaction: synergy, FICI≤0.5; antagonism FICI >4.0; and no interaction FICI from 0.5 to 4.0.

Post-antibiotic effect assays. Exponential grown cultures (OD_600_= 0.5-1.0) were diluted to a final cell density of 1.0×10^5^ cells/mL in 10 mL supplemented 7H9-based broth medium (i.e., total 1×10^6^ cells) and treated with the compounds for 2 hours in roller bottles (5 rpm) at 37°C. After the treatment period, compounds were washed out by three centrifugation steps (3,500 rpm), supernatants removed and cells resuspended in 1 mL of fresh supplemented 7H9-based broth medium plus tyloxapol (final cell density 10^6^ cells/mL). CFU counting was performed for every sample to determine whether cell killing has occurred during the 2 hours of treatment. From this cell suspension, 100 µL were then added to BD BBL^TM^ Mycobacteria Growth Indicator tubes (MGIT^TM^) (total of 1.0×10^5^ cells/tube) in duplicates and the Time-To-Positivity (TTP, hours) measured in a BACTEC MGIT instrument. Positivity was defined as a Growth Index higher than 75 (GI > 75).

Time kill assay. Frozen stocks of M. tuberculosis H37Rv were inoculated in 7H9-based broth supplemented with glycerol and ADC, without tyloxapol. Cultures were incubated at 37°C for three days to allow for bacterial recovery and exponential growth phase. Then 10-mL were inoculated in 25 cm^2^ tissue culture flasks (or roller bottles) to a typical final cell density of 10^5^ cells/mL and drugs added at the designated times, concentrations and combinations depending on the type of assay, i.e., dose-response or replenishment. At every time point, cultures were thoroughly mixed, samples (100 µL) 10-fold serially dilute in 1x PBS buffer with 0.1% tyloxapol and 100 µL plated on 7H10 agar plates supplemented with 10% OADC. Agar plates were then processed as described above.

MIC determination against M. tuberculosis clinical isolates. The Minimum Inhibitory Concentration (MIC) of each tested compound against M. tuberculosis clinical isolates was determined in 96-well flat-bottom, polystyrene microtiter plates in Middlebrook 7H9 broth base supplemented with 10% ADC and 0.05% tyloxapol. Ten two-fold serial dilutions of the test compounds dissolved in DMSO were performed from column 1 to 10. Moxifloxacin was used as a dose response compound control starting at 1 μg/mL in column 11. Column 12 was used for positive (cells, no drug) and negative (no cells, no drug) controls. Similar plates with the same layout were also prepared including in the 7H9 medium 4-6 μg/mL of potassium clavulanate in order to test the potential MIC shift in the presence of this β-lactamase inhibitor. M. tuberculosis cells were grown to exponential phase (ca. OD_600_= 0.2-0.3) and standardized to 1×10^7^ CFU/mL (ca. OD_600_= 0.125). The pre-inoculum was then further diluted (1:200) in the assay medium and 200 µL added to each well (final cell density ca. 5×10^4^ CFU/mL). Plates were then incubated in sealed boxes to prevent drying of the perimetric wells and incubated without shaking at 37°C for 6 days (REMA readout) or 2 weeks (visual inspection readout). For the REMA readout [22], a resazurin solution was prepared by dissolving one tablet of Resazurin (Resazurin Tablets for Milk Testing; Ref. 330884Y VWR International Ltd) in 30 mL of sterile PBS (phosphate buffered saline). Of this solution, 25 μL were added to each well and the fluorescence measured after 48 hours in a Spectramax M5 (Molecular Devices) reader (λ_Ex_= 530 nm, λ_Ex_= 590 nm, cut-off 570 nm) to determine the MIC value. For the visual inspection readout, at weeks 1 and 2, plates were read with inverted enlarging mirror plate reader and individual wells grade as either growth or no growth by visual inspection. Regardless of the readout method used, the MIC value was defined as the lowest concentration that inhibited growth compared to untreated controls. The MIC_90_ value was considered the concentration that inhibited the growth of 90% of the clinical isolates.

Single-cell time-lapse microscopy. Experiments were performed as previously described [23]. Briefly, sanfetrinem was always prepared fresh in water solution (5 mg/mL), filter-sterilized and added to 50 mL of 7H9 broth at tests concentrations. A M. bovis BCG-gfp primary culture was started from a frozen stock in 10 mL of complete 7H9 supplemented with 50 µg/mL of Hygromycin B and allow to grow at 37°C in a 25 cm^2^ tissue culture flask until mid-exponential phase (OD_600_= ca. 0.5). To prepare a single cell suspension, cells (2×1 mL) were pelleted, resuspended in 200 µL of complete 7H9 without tyloxapol and hygromycin B, and passed through a 5 µm syringe filter. This suspension (4 µL) was seeded onto a microfluidic device, which was assembled and transported to the microscope (Deltavision Elite Imaging System, Ref: 53-852339-002) for imaging capture. For every experiment, 110 points were labeled and acquired every 4 hours for 15 days using two microscopy channels: contrast phase (Pol, Blank, 32%T 0.01 seg) and fluorescence (GFP, 100%T 0.01 seg). A CoolSnap HQ camera was used with a 10x gain, 100x objective and autofocus system on contrast phase channel (7 µm on Z axis/0.2 µm steps). Sanfetrinem was infused into the microfluidic devise with a medium flow rate of 20 µL/min. Sanfetrinem-containing 7H9 media solution was refreshed every 24 hours to account for thermal drug degradation at room temperature; as such, we can assume that sanfetrinem concentration was maintained constant throughout the experiments. After a 72 hours growth phase, sanfetrinem was infused at a constant concentration of 20 μM (6xMIC; MIC_M. bovis BCG_= 3.2 μM) and maintained for 144 hours (6 days), after which, sanfetrinem pressure was removed and cells allowed to recover for 3 additional days; a staining solution of propidium iodide (PI) was then added to differentiate death/alive cells.

*in vivo* assays in murine models of M. tuberculosis infection. The *in vivo* anti-tubercular activity of sanfetrinem, and other β-lactams, was evaluated following an established experimental design using DHP-1 knockout (Dpep-1 KO) mice in 129sv background, instead of the previously described C57Bl/6 background [24]. In brief, pathogen-free, 8-10 weeks-old female 129sv DHP-1 KO mice were purchased from either Envigo or Taconic DK and allowed to acclimate for one week. Mice were intra-tracheal infected with approximately 10^5^ CFU/mouse (M. tuberculosis H37Rv). Treatment was started nine days after infection and administered twice a day (bid) from day 9 to day 14 after infection at the following doses [mg/kg bw]: amoxicillin [200], clavulanate [100], faropenem [500], meropenem [300], sanfetrinem sodium salt [200] and sanfetrinem cilexetil [400] (it is estimated that a dose of sanfetrinem cilexetil 400 mg/kg bw equals to a dose of sanfetrinem 250 mg/kg bw). Meropenem and sanfetrinem sodium salt were administered subcutaneously while all other β-lactams were administered by oral gavage. Lungs were harvested on days 9 and 15 and lung lobes aseptically removed, homogenized and frozen. Homogenates were thaw, plated in 10% OADC-7H11 medium + 0.4% activated charcoal and CFU enumerated after 18 days of incubation at 37°C. Differences in bacterial lung burden [log_10_ CFU/mouse (lungs)] of treated mice versus untreated controls (Day 15 after infection) were analyzed by the one factor ANOVA, Dunnet’s posttest and Bonferroni’s multiple comparisons tests using GraphPad Prism 7.0. For the pharmacokinetic studies of sanfetrinem and sanfetrinem cilexetil, animals were dosed orally at 200 mg/kg and 400 mg/kg, respectively, and blood samples taken and analyzed at 10, 60, 180 and 360 minutes post dosing. All animal studies were ethically reviewed and carried out in accordance with European Directive 2010/63/EU and the GSK Policy on the Care, Welfare and Treatment of Animals.

Drug quantification. Ultra-Performance Liquid Chromatography-Mass Spectrometry/Mass Spectrometry Analysis (UPLC-MS/MS) was used to quantify the amount of sanfetrinem, in 7H9 medium and blood samples. The UPLC-MS/MS system consisted of an Acquity UPLC series (Waters Corporation, Madison, USA) coupled with a Sciex API 4000 instrument (AB Sciex, Toronto, Canada). Fifteen microliters of every sample were added to 190 µL of protein precipitant buffer (acetonitrile /methanol 80:20 v/v containing the internal standard) and filtered through a 0.45 µm pore size filter. From each sample, 150 µL were evaporated and re-suspended with 150 µL of milli-Q water before LC-MS/MS analysis. Samples were then loaded into an Acquity UPLC HSS T3 50 x 2.1mm, 1.8 µm column (Waters Corporation, Madison, USA) and eluted by gradient with 10 mM Ammonium Formate plus 0.1% Formic Acid and Acetonitrile at a flow rate of 0.4 mL/min. The MS/MS system was operated in MRM negative mode with 280.1/236.3 transition.

## RESULTS

An intracellular screening of β-lactams identifies the anti-mycobacterial activity of sanfetrinem. A GSK in-house library of 1,973 unique β-lactams was single-shot (50 µM) screened for activity against M. tuberculosis in infected THP1 macrophage cells, and 105 active hits were identified with ca. 70% inhibition cutoff (Figure 1A). Forty-five compounds were confirmed active in successive, more stringent intracellular dose-response studies (90% inhibition cut-off), yielding a hit discovery rate of 2.28% (Table S1). Dose response studies were also performed under extracellular (7H9) conditions and IC_90_ values compared versus the intracellular (THP1) activity. Three main classes of β-lactams were identified based on their activity profile (cut-off 20 µM): active only intracellular, active only extracellular or with dual activity. Sanfetrinem cilexetil (GV118819) was identified with dual activity (Figure 1B), which was also confirmed for the parent sanfetrinem compound (GV104326). Interestingly, both displayed slightly increased activity under intracellular conditions compared to extracellular conditions (Figure 1C). The intracellular activity of sanfetrinem was comparable to that of the known intracellularly active cefdinir [25], and higher than other clinically approved β-lactams (Figure S1).

**Figure 1.**
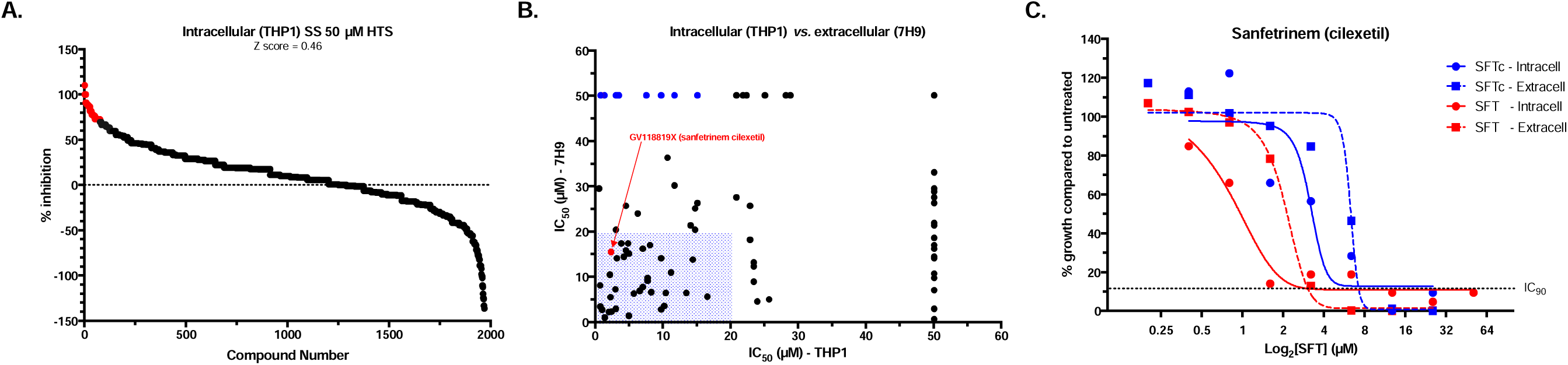
Identification of sanfetrinem in a β-lactam intracellular screening assay. A library of 1,973 unique β-lactams was screened for activity against M. tuberculosis in infected THP1 macrophage cells. (A) Single shot (50 µM) assays identified 105 hits with at least 70% intracellular growth inhibition compared to untreated (red dots); (B) Dose response studies calculated the IC_50_ values of the 100 selected hits in both intracellular (THP1) and extracellular (7H9) conditions. Some β-lactam hits displayed exclusive intracellular activity (blue dots) while sanfetrinem cilexetil (red dot) displayed dual activity (cutoff 20 µM, dashed blue box); (C) Sanfetrinem cilexetil (SFTc) and sanfetrinem (SFT) in intracellular (THP1 cells) and extracellular (7H9 medium) dose response studies.

Media composition impacts the activity of sanfetrinem against M. tuberculosis. Dose response studies were performed under different media conditions to assess their impact on sanfetrinem’s activity, i.e., standard media including dextrose and glycerol vs. glucose vs. cholesterol, detergents typically used in the assay medium, bovine serum albumin (BSA) and the presence of β-lactamase inhibitors (Figure 2). Cholesterol is one of the main carbon sources available to mycobacteria at the site of infection [26]. To test the activity of sanfetrinem with cholesterol as the only carbon source the detergent tyloxapol needs to be added in order to solubilize it into the assay media, which is known to alter the cell wall structure and improve the activity of some cell wall targeting compounds, such as ethambutol [27]. In fact, the activity was improved in the presence of tyloxapol for some of the β-lactams used in this study, i.e., cefadroxil, cefdinir and faropenem, but not for amoxicillin or meropenem (Figure S2). Similarly, the activity of sanfetrinem was not affected by the presence of tyloxapol, which provides confidence in the observed improved activity (4-fold) in the presence of cholesterol as the only carbon source, a more physiologically relevant condition than the standard broth media supplemented with dextrose and glycerol (Figure 2A) or simply supplemented with glucose (Figure 2B). In this case, the presence of BSA in the assay media had a significant impact, decreasing sanfetrinem’s activity 4-fold (Figure 2B), thus suggesting protein binding as a key parameter to be considered for efficacy projections in pre-clinical and clinical development. Finally, the addition of clavulanate to the assay medium had a minor impact on sanfetrinem’s activity against the laboratory strain M. tuberculosis H37Rv and a slight 2-fold improvement in a rifampicin resistant (H526D) M. tuberculosis H37Rv strain (Figure 2C), although with this set of experiments the actual contribution of clavulanic acid to the *in vitro* activity of sanfetrinem could not be fully discerned.

**Figure 2.**
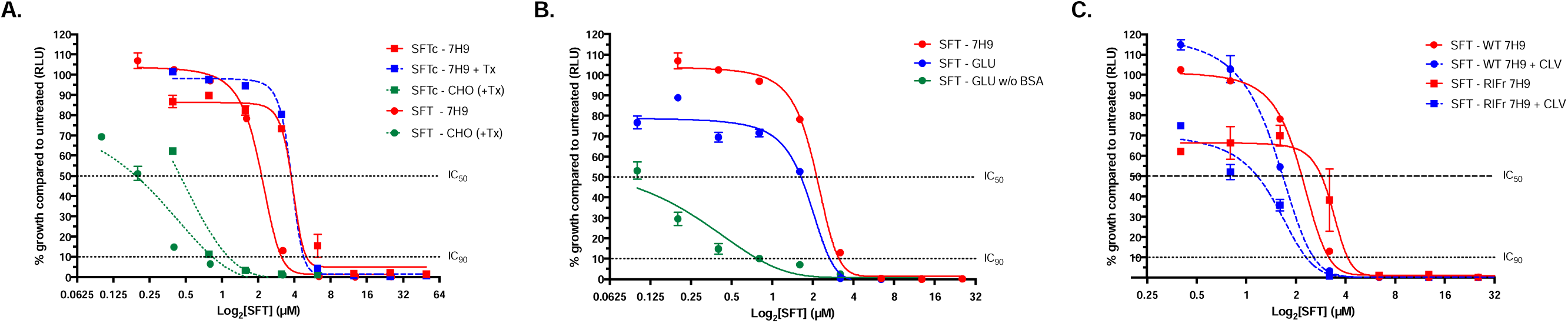
*in vitro* characterization of sanfetrinem against M. tuberculosis. Dose response studies of sanfetrinem in (A) standard 7H9 medium (7H9), 7H9 plus the detergent tyloxapol (7H9 + Tx), 7H9 salts containing cholesterol as the only carbon source (CHO+Tx); (B) 7H9 salts containing glucose as the only carbon source with (GLU) and without BSA (GLU w/o BSA) and; (C) standard 7H9 medium plus the addition of clavulanic acid (7H9 + CLV). SFTc, sanfetrinem cilexetil; SFT, sanfetrinem; WT, wild-type M. tuberculosis H37Rv strain; RIFr, rifampicin-resistant M. tuberculosis H37Rv H526D strain. Unless otherwise specified, experiments were performed with the wild-type strain.

Broad spectrum activity of sanfetrinem against M. tuberculosis clinical isolates. In order to assess the potential for global implementation of sanfetrinem as a new anti-TB drug and the role clavulanate might play in a combined therapy, its anti-tuberculosis activity was tested against a panel of M. tuberculosis strains (n= 61), including drug-susceptible, MDR and XDR clinical isolates from different geographical locations alone and in the presence of clavulanate. The activity of sanfetrinem was compared to that of the clinically proven meropenem, among other clinically used β-lactams. Sanfetrinem was the most active β-lactam with an MIC_90_ value of 2-4 µg/mL, while meropenem displayed an MIC_90_ of 4-16 µg/mL, with an up to 16-fold increase in activity in the presence of clavulanate. Amoxicillin was less active (MIC_90 (AMX)_ > 32 µg/mL); however, its activity was strongly enhanced up to 32-fold in the presence of clavulanate (MIC_90 (AMX/CLV)_ = 2-8 µg/mL). Sanfetrinem showed a 4-fold increased activity in the presence of clavulanate (MIC_90 (SFT/CLV)_ = 0.5-2 µg/mL) (Figure 3). As in the dose response studies with the rifampicin-resistant H37Rv strain (Figure 2C), a detailed analysis revealed that strains with a resistant profile were more likely to benefit from the addition of clavulanate to sanfetrinem therapy (Table S2); kill kinetic assays with a rifampicin resistant strain confirmed this observation (Figure S3). In summary, sanfetrinem displayed the best *in vitro* activity of all β-lactams tested against M. tuberculosis clinical isolates with strong potential for broad clinical coverage, i.e., active against DS, MDR and XDR strains. Concomitant treatment with clavulanic acid might improve the effectiveness of sanfetrinem.

**Figure 3.**
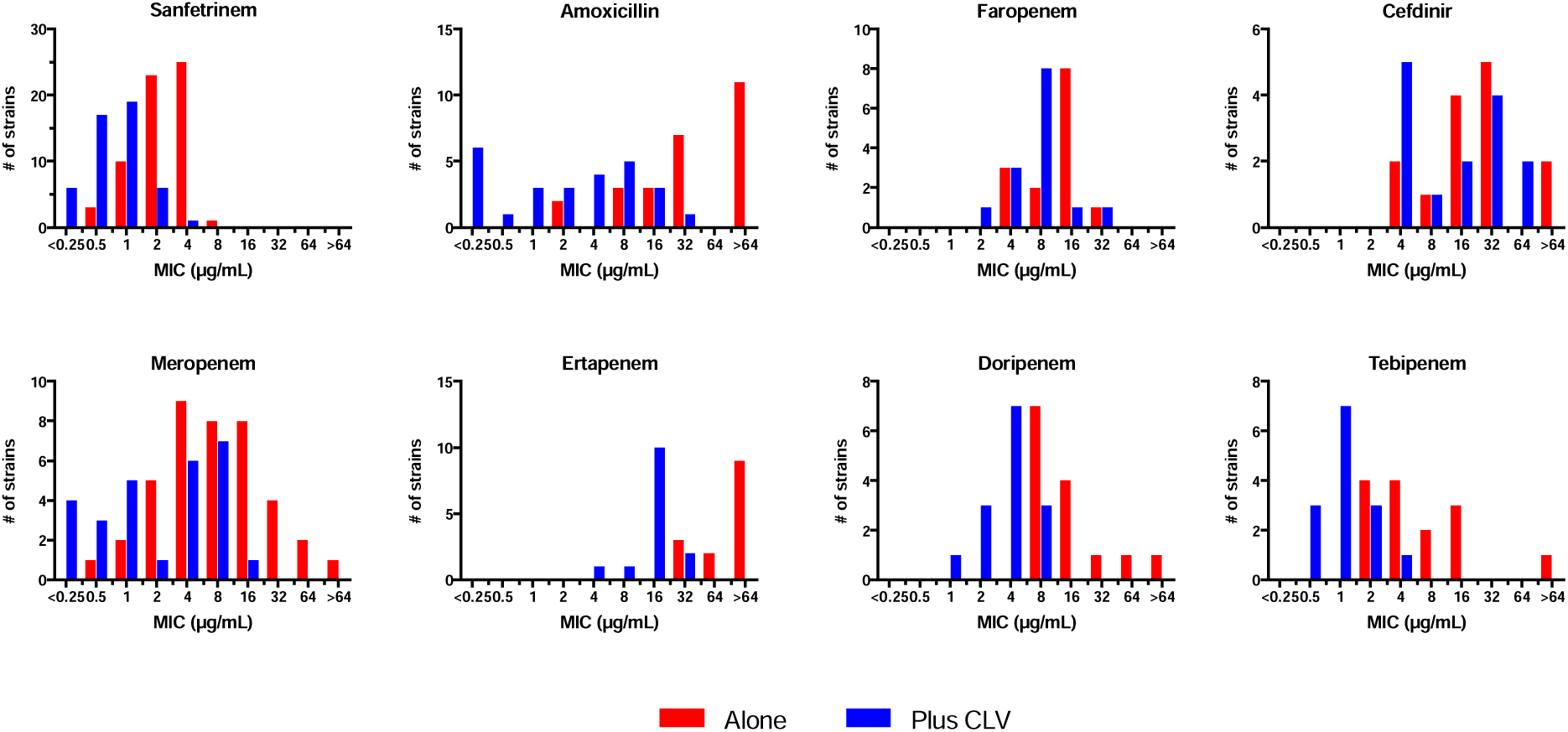
Antimicrobial *in vitro* activity of sanfetrinem and other β-lactams against M. tuberculosis clinical isolates. The activity of several β-lactams was tested against a panel of M. tuberculosis drug-susceptible, multi-drug resistant and extensively drug resistant clinical isolates, together with laboratory strains alone and in combination with clavulanic acid (Plus CLV). The number of strains tested for the compound alone or in the presence of clavulanic acid were, respectively: sanfetrinem (62/49), meropenem (40/27), amoxicillin (26/26) and tebipenem, ertapenem, doripenem, faropenem and cefdinir (14/14). Detailed MIC values and resistant profiles of the clinical isolates are described in Table S2. CLV, clavulanic acid.

Sanfetrinem displays rapid bactericidal *in vitro* activity against M. tuberculosis. Having identified sanfetrinem in an intracellular single-shot screening assay (Figure 1); CFU-based intracellular time-kill assay confirmed these data: β-lactams are active against actively replicating bacteria and highly unstable in the assay media; thus, sanfetrinem was administered twice (after 1- and 4-days post-infection) to capture the phase of active bacterial replication. A modest bacteriostatic activity could be observed at the highest concentration tested (50 µM) when sanfetrinem was just added once at day 1 post-infection. However, a second dose of sanfetrinem after 4 days of infection (presumably when bacteria have adapted to intracellular growth and are actively replicating) showed a significant effect with bactericidal activity even at a low 1.25 µM concentration (Figure 4A). After validating the intracellular bactericidal activity of sanfetrinem against M. tuberculosis, we aimed to further characterize its activity in extracellular conditions. We first performed stability assays of sanfetrinem in the assay media and calculated a half-life (T_1/2_) of ca. 0.6 days, which effectively means that no drug is present in the assay media after 4 days of incubation. Regardless of this degradation, dose response time-kill assays showed a rapid bactericidal activity over the first 24 hours of exposure at concentrations as low as 5 µM. At higher concentrations, this killing was maintained for up to seven days, even reaching the limit of detection with the highest concentration tested. In all conditions, bacterial regrowth was observed after the initial bacterial killing (Figure 4B). Colonies isolated from the regrowth showed similar MIC values as the wild-type, suggesting this was due to the lack of drug in the assay medium and not to the acquisition of genetic mutations. The time to regrowth correlated with the initial amount of sanfetrinem, as confirmed by time-to-positivity (TPP) studies (Figure 4C). As such, and considering the T_1/2_ of sanfetrinem, we performed top-up time-kill assays at a fixed 5 µM concentration. Sanfetrinem was replenished in the assay media every day during 3 or 6 consecutive days. As shown in Figure 4D, time to regrowth was dependent on maintained exposure levels of sanfetrinem. Importantly, when drug exposure was maintained, sanfetrinem was also bactericidal against a high inoculum (10^7^ cell/mL). Nevertheless, even after one single dose addition of sanfetrinem, there was a lag-phase to regrowth of almost 7 days. This result could suggest a post-antibiotic effect of sanfetrinem (Figure 4E), an *in vitro* feature not observed for other β-lactams (Figure S5).

**Figure 4.**
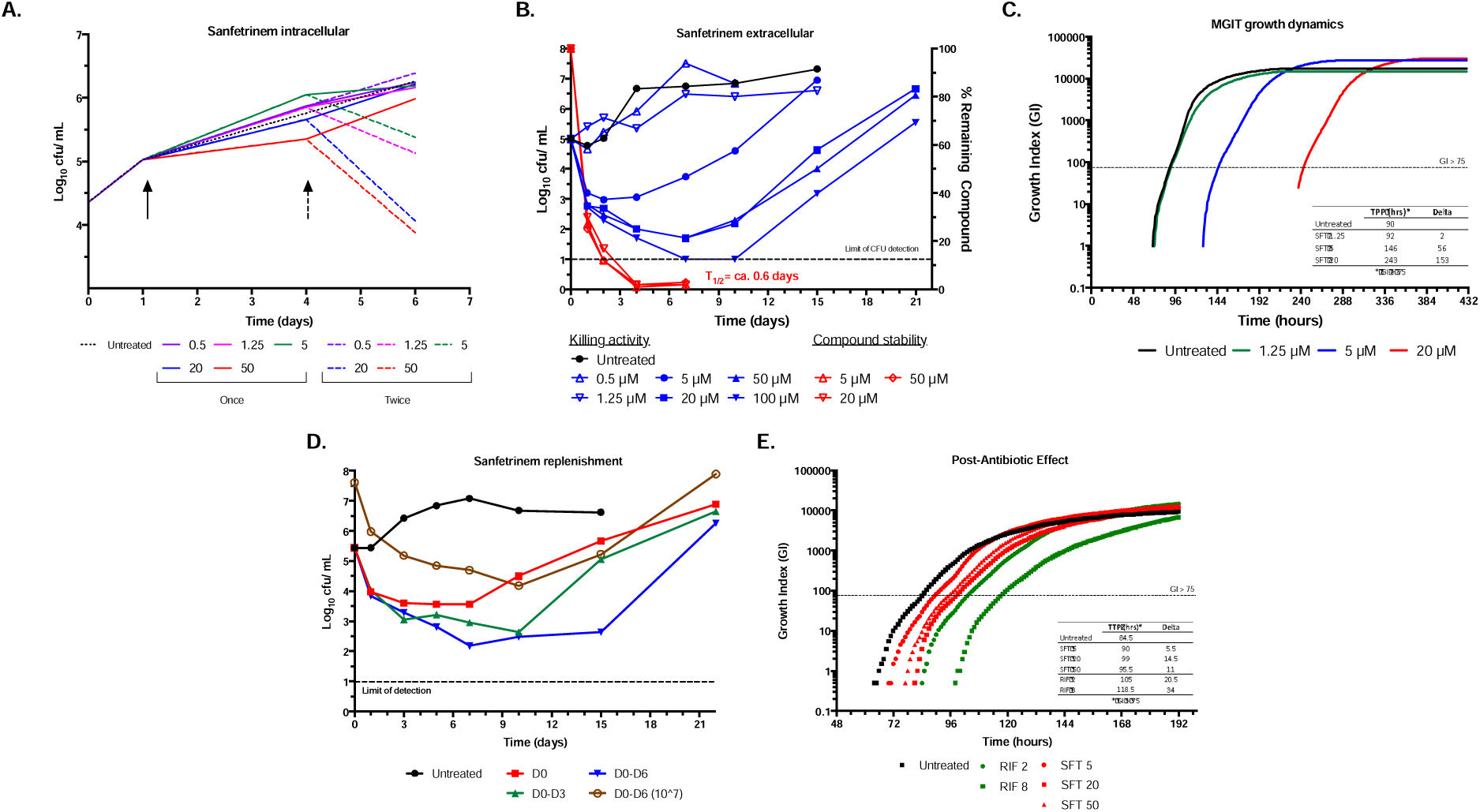
Intracellular and extracellular time-kill and post-antibiotic effect assays of sanfetrinem against M. tuberculosis. (A) CFU-based intracellular dose response assays were performed with sanfetrinem (µM) added only on day 1 (solid arrow) or twice on days 1 and 4 (dashed arrow). (B) Dose response time-kill assay of sanfetrinem in standard 7H9 medium. Blue lines indicate bacterial density over time after sanfetrinem treatment. Red lines indicate the amount of sanfetrinem present in the assay media due to thermal instability at 37°C. (C) The activity of sanfetrinem against a similar M. tuberculosis as in time-kill assays was continuously monitored in the MGIT system and the time-to-positivity (TPP) calculated. Delta indicates the elapsed time to reach positivity (GI > 75) compared to the untreated sample. (D) Sanfetrinem was added at 5 µM only once at day 0, or daily from day 0 to day 3 or from day 0 to day 6 to replenish to the target concentration, accounting for a degradation T_1/2_ of ca. 0.6 days. Two starting inocula were also used: approx. 10^5^ and 10^7^ cells/mL. Cell density (CFU/mL) was followed over time. (E) The post-antibiotic effect of sanfetrinem was measured against M. tuberculosis H37Rv. Cells treated at the corresponding drug concentrations for 2 hours were transferred (10^5^ cells) to MGIT tubes and the TTP measured. The 2-hours treatment did not affect CFU counts. Rifampicin was included as positive control. Numbers in legend indicate drug concentration in µg/mL for rifampicin and in µM for sanfetrinem (5 µM equals to ca. 1.7 µg/mL). Delta indicates the increase in TTP compared to untreated. RIF, rifampicin; SFT, sanfetrinem.

To further characterize the activity of sanfetrinem at the single cell level we performed time-lapse microscopy studies (Figure 5). Unlike the time-kill assays, the microfluidics device allowed us to maintain the concentration of sanfetrinem over the treatment period. A rapid killing profile due to abrupt cell lysis was observed only 8 hours after drug exposure from which cells were not able to recover after removing sanfetrinem six days later (Figure 5A). Remaining bacteria exposed to sanfetrinem were PI positive which indicates a compromised cell membrane with a distorted cell morphology (Figure 5B). All together, these studies demonstrated the rapid bactericidal activity of sanfetrinem against Mycobacterium in both extracellular and intracellular conditions, with a long-lasting effect when drug pressure was maintained over the course of the experiment.

**Figure 5.**
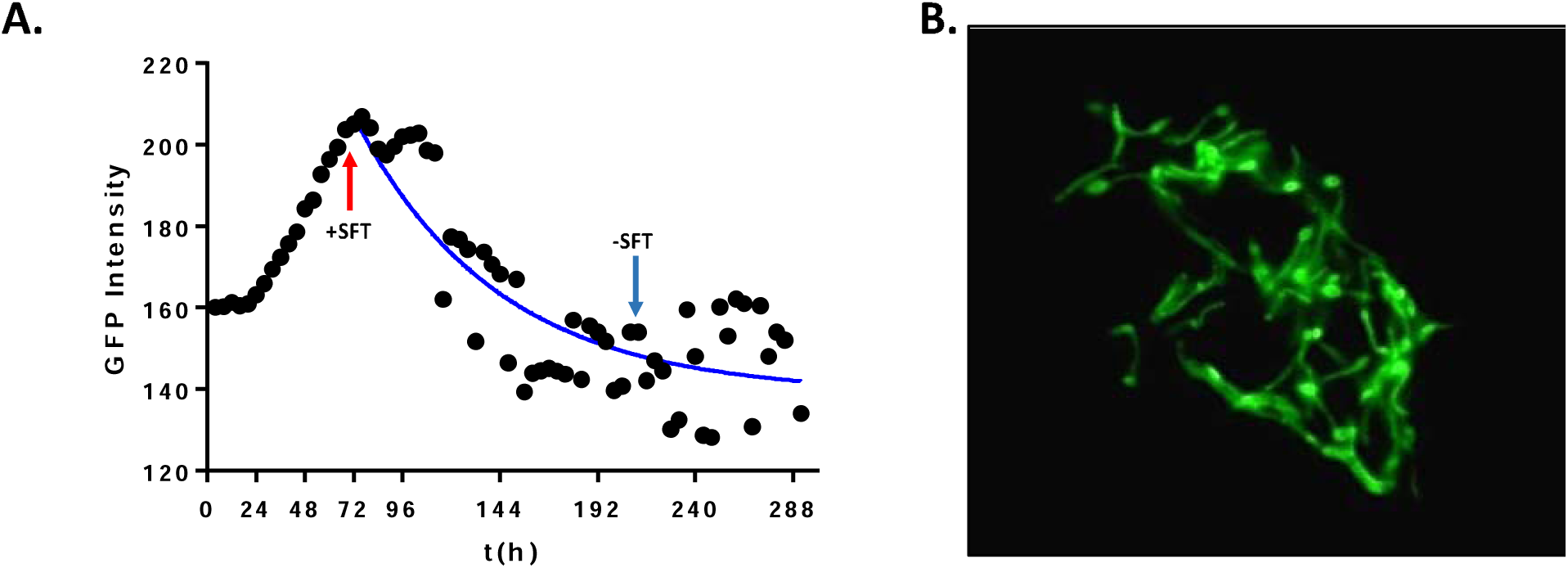
Time-lapse microscopy of the effect of sanfetrinem against M. bovis BCG. (A) M. bovis BCG cells were allowed to adapt for 72 hours until exponential growth was achieved in the microfluidic device. After 72 hours, sanfetrinem was added (+SFT) at a fixed concentration of 20 µM for 6 days. Then, sanfetrinem was removed (-SFT) and bacterial cells allowed to recover for 3 days before propidium iodide staining. (B) Bacterial morphology after 6 days exposure to sanfetrinem.

Sanfetrinem displays strong *in vitro* synergistic interactions with amoxicillin, ethambutol, rifampicin and rifapentine. *in vitro* synergy assays were performed with sanfetrinem and meropenem in combination with a panel of twenty-six clinically approved (or with known mode of action) antimicrobials to further explore the repurposing potential of sanfetrinem within a TB regimen (Figure 6). Sanfetrinem displayed strong synergy with amoxicillin and rifampicin with FICI values lower than 0.25 and up to 8-fold reductions in MIC values within the combination, compared to the MIC of the compound alone; additionally, synergy was also observed with rifapentine (FICI = 0.38) and ethambutol (FICI = 0.5). This pattern of interaction differed from that of meropenem for which synergy was more prevalent, although strong synergy (FICI<0.25) was only observed for rifampicin, rifapentine and cephradine, but not for amoxicillin (Figure 6A). A closer look at the sanfetrinem synergistic combinations revealed a shift in the dose response curves when in the presence of sub-MIC concentrations of the companion drugs (Figure 6B), consistent with the observed FICI values. The observation that sanfetrinem and meropenem had different synergy profiles was not uncommon as it has been observed for other β-lactams [19].

**Figure 6.**
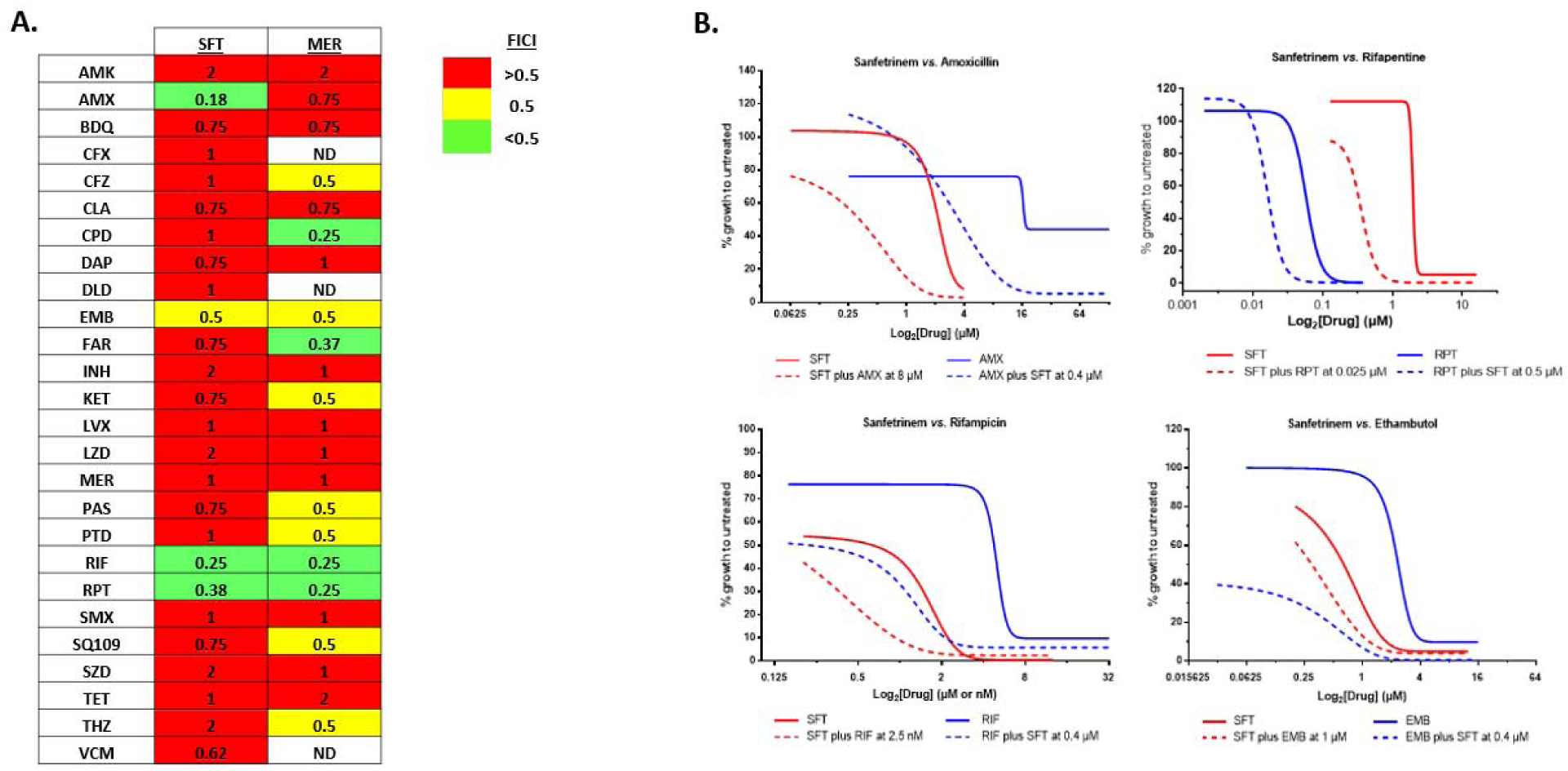
*in vitro* synergy profiling of sanfetrinem and meropenem against M. tuberculosis H37Rv. (A) Fractional inhibitory concentration indexed (FICI) of sanfetrinem and meropenem in pair-wise combinations of clinically approved or with known mode of action antimicrobials (n= 26). A FICI ≤0.5 indicates synergy, while a FICI >0.5 and ≤4 indicates no interaction; (B) Dose response interaction curves of sanfetrinem (red) with synergistic partners (blue). Solid lines represent the dose response curve of drugs alone while broken lines represent the dose response curves in the presence of sub-MIC concentrations of the synergistic partner. Concentrations of drugs are expressed in µM, except for rifampicin and rifapentine that are expressed in nM. AMK, amikacin; AMX, amoxicillin; BDQ, bedaquiline; CFZ, clofazimine, CLA, clarithromycin; CPD, cephradine; DAP, dapsone; DLD, delamanid; EMB, ethambutol; FAR, faropenem, INH, isoniazid; KET, ketoconazole; LVX, levofloxacin; LZD, linezolid; MER, meropenem; PAS, p-amino salicilic acid; PTD, pretomanid; RIF, rifampicin; RPT, rifapentine; SFT, sanfetrinem; SMX, sulfamethoxazole; SZD, sutezolid; TET, tetracycline; THZ, thioacetazone; VCM, vancomycin.

Time-kill assays were performed to study the *in vitro* pharmacodynamics of sanfetrinem combinations against M. tuberculosis. The activity of sanfetrinem, amoxicillin/clavulanate, and rifampicin was evaluated through the examination of their individual and combined effects with the compounds under investigation, I.e., alone and in pair-wise and triple combinations. Drugs were added at sub-optimal concentrations in order to allow detection of synergistic interactions with an increased killing capacity, demonstrated by higher CFU decline of the combination compared to the drugs alone. As shown in Figure 7, at the tested concentration sanfetrinem was able to reduce the bacterial burden by 2-logs over the first 24 hours, but regrowth occurred after 2 days, similar to previous observations (Figure 4); this bactericidal effect was stronger than that observed for amoxicillin/clavulanate (with only 1-log reduction over the first 24 hours). The combination of sanfetrinem with amoxicillin/clavulanate exhibited a minor interaction characterized by a slight delay in re-growth. However, the combination of either sanfetrinem or amoxicillin/clavulanate with rifampicin had a strong positive interaction with a post-antibiotic effect between 10-15 days preventing the culture from resuming growth. Triple combinations of sanfetrinem and amoxicillin/clavulanate plus either rifampicin had profiles similar or slightly better that the respective pair-wise combos. In both cases, a sanfetrinem-dependent rapid initial killing was observed in addition to maintaining bacteriostasis up to 15 days after treatment (Figure 7). Our studies thus confirm rifampicin and rifapentine as strong synergistic partners of sanfetrinem and amoxicillin/clavulanate with the potential to be part of a novel combination therapy.

**Figure 7.**
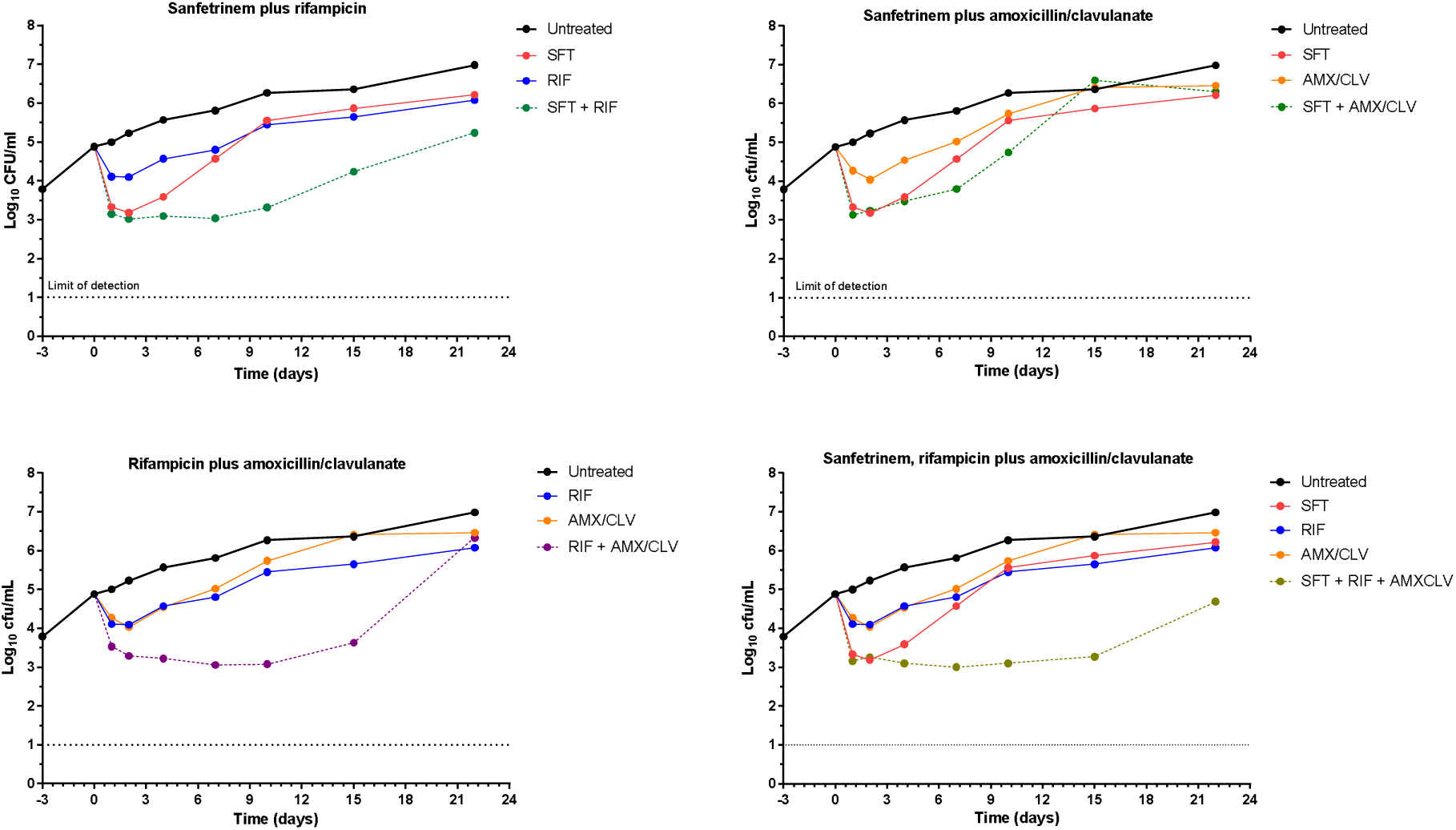
Combinatorial time-kill assays of sanfetrinem with rifampicin and amoxicillin/clavulanate. Sanfetrinem (SFT), rifampicin (RIF), and amoxicillin/clavulanate (AMX/CLV) were tested alone and in pair-wise and triple combinations against M. tuberculosis H37Rv in time-kill assays. Cultures were allowed to adapt for three days to exponential growth conditions (Day-3). Then, drugs were added at day 0. Concentrations used were: SFT = 2.5 µM (0.87 µg/mL); RIF = 0.5 µM (0.41 µg/mL); AMX = 2.2 µM (0.8 µg/mL); CLV = 5 µg/mL.

Sanfetrinem is active against M. tuberculosis in an *in vivo* model of infection. β-lactams display poor *in vivo* efficacy in traditional murine models of M. tuberculosis infection [28]; [29]; [30]. One of the reasons for this lack of efficacy is the increased expression of the murine renal dehydropeptidase (DHP-1) enzyme, several orders of magnitude higher than its human homologue, which limits the exposure of β-lactams in such models. However, developments in the field have demonstrated robust and significant CFU reductions (1 to 2 log CFU) in the lungs of infected DHP-I-deficient mice, in comparison to the untreated control [24].

In this study, we used DHP-1 knockout mice to evaluate the *in vivo* activity of sanfetrinem in comparison with β-lactams already tested in clinical trials (Figure 8). A first experiment compared the activity of sanfetrinem cilexetil with a combination of faropenem plus clavulanate. Untreated controls experienced an increase in lung CFU of ca. 1-log over the treatment period (day 9 vs. day 15), while treatment groups were able to prevent such growth. In a second experiment, the activities of both forms of sanfetrinem (sodium salt dosed subcutaneously and cilexetil dosed orally) were compared against combinations of meropenem plus either clavulanate or clavulanate and amoxicillin. Similarly, untreated controls displayed an increase in lung CFU over the treatment period (ca. 1.6-log CFU) while this growth was prevented in all treatment groups. When comparing among treatment groups, the combination of meropenem-clavulanate-amoxicillin was the most active, although not significantly different from the sanfetrinem and meropenem-clavulanate groups. The oral sanfetrinem cilexetil arm also resulted in a significant lung CFU reduction, although slightly less than the most active meropenem-clavulanate-amoxicillin group. This could be due to the lower exposure levels obtained with the oral form of sanfetrinem compared to the subcutaneous administration, as observed in mice pharmacokinetic studies (Figure S5). However, no significant differences were observed between the activities of the two forms of sanfetrinem.

**Figure 8.**
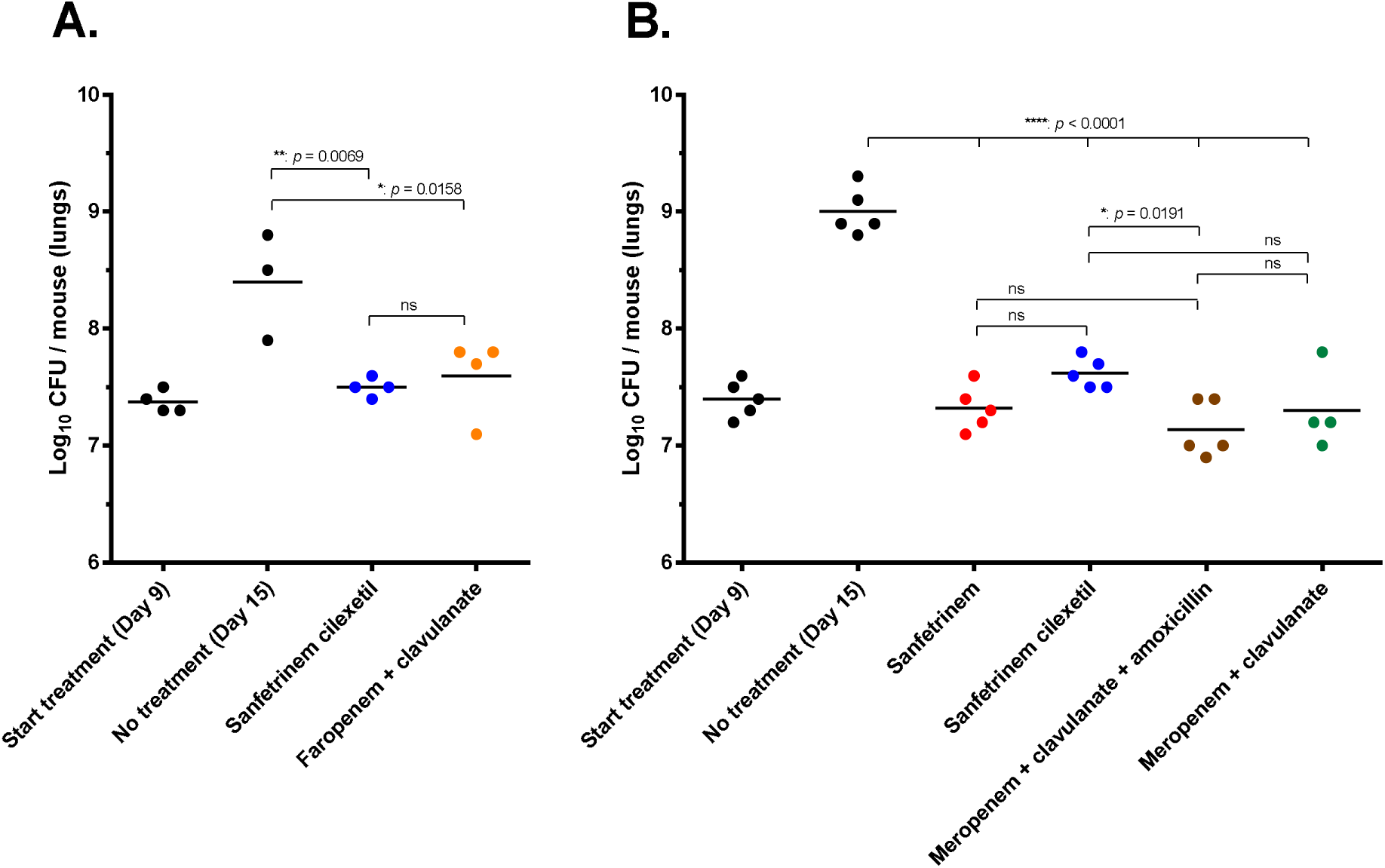
Anti-tubercular activity of different β-lactams in an acute murine model of M. tuberculosis infection. Two independent experiments (A and B) were performed in which each point represents a data value from a single DHP-1 knockout mouse. Treatment was started nine days after infection and administered twice a day (bid) from day 9 to day 14 after infection. Lungs were harvested on day 15. Meropenem and sanfetrinem were administered subcutaneously and sanfetrinem cilexetil and all other β-lactams by oral gavage at the following doses (mg/kg bw): amoxicillin (200), faropenem (500), meropenem (300), sanfetrinem sodium salt (200), sanfetrinem cilexetl (400) and clavulanate (100). ANOVA analysis and Dunnet ś posttest versus the “No treatment (Day 15)” group indicates significant differences upon treatment for all groups (p-value <0.05). Bonferroni’s multiple comparisons tests show significant differences among treatment groups. ns, no significant.

## DISCUSSION

A seminal Phase 2a EBA clinical study demonstrated the efficacy of meropenem in combination with amoxicillin/clavulanate for the treatment of TB [12]. Subsequent studies in treating MDR/XDR patients supported these observations [31], although recent studies with drug-susceptible TB cast some doubts on the role of meropenem in programmatic TB treatment due to poor tolerability [32]. In addition, due to the lengthy therapies required for TB treatment, the clinical use of meropenem comes with some limitations, mainly related to the intravenous route of administration and its susceptibility to degradation by intrinsic *Mtb* β-lactamases. The stated goals of this screening of β-lactams, and related ones [3] against *Mtb* were to select compounds with: (i) similar or greater potency as compared to the clinically proven meropenem; (ii) inherent stability to BlaC degradation; and (iii) oral bio<u>a</u>vailability. Therefore, we were excited and pleased to identify sanfetrinem cilexetil, a tricyclic β-lactam candidate, met such criteria.

Tricyclic carbapenems or “trinems” were originally reported by Glaxo Wellcome in 1966 [14, 33] with the original aim of inhibiting β-lactamases [34],[35]. Several *in vitro* and *in vivo* studies demonstrated the efficacy of sanfetrinem against Gram-positive and Gram-negative bacteria [36](Unpublished Data). The oral prodrug sanfetrinem cilexetil was subsequently developed for respiratory tract infections in adults and otitis media in children. Multiple successful Phase 2 trials in these indications demonstrated its ability to achieve therapeutically relevant exposures upon oral dosing. With just a slightly increased tendency toward gastrointestinal tolerability issues in patients taking sanfetrinem cilexetil, these trials and preceding development showed a reasonable safety profile, in line with other β-lactam drugs, including pediatric populations (GSK, unpublished data). Compatibility with pediatric and pregnant populations is a major gap in TB drug development, representing another exciting feature for sanfetrinem cilexetil.

β-lactams are rapid bactericidal antibiotics that were largely considered inactive against intracellular pathogens [37]. *Mtb* can be found extracellularly in the caseum of the granulomas but mainly resides in intracellular compartments at the site of infection. Effectively targeting both populations will contribute to TB treatment shortening and control of the disease by means of a rapid decline in extracellular burden and reaching difficult-to-treat intracellular bacteria. We thus designed an intracellular screening program to test a GSK in-house library of 1,973 unique β-lactams; sanfetrinem cilexetil was identified as one of the most promising hits from this screen with dual intracellular and extracellular activity (Figure 1). This finding is consistent with previous reports showing entry and intracellular accumulation of sanfetrinem into human polymorphonuclear granulocytes [38]. Time-kill and time-lapse microscopy assays further demonstrated the rapid bactericidal capacity of sanfetrinem in both intracellular and extracellular conditions and an extended lag phase despite the poor chemical stability under assay conditions (Figure 4 and Figure 5). *Mtb in vitro* assays require several weeks to produce results, and compound stability experiments showed that sanfetrinem (like other carbapenems [7] is almost entirely degraded within 1-2 days. Quantifying the actual carbapenem’s activity in *Mtb in vitro* assays is non-obvious, with extensive effort and coordination needed due to the biohazard nature of the samples containing a Biosafety Security Level 3 pathogen. The implication of this instability is that traditional MIC determinations (typically used as the main initial proxy for *in vitro* activity) for carbapenems are misleading because their antimicrobial activity is consistently underestimated due to the drug being present only for a fraction of the time assay [39]. Time-kill replenishment studies accounting for the drug degradation in the assay media demonstrated a better proxy to quantify sanfetrinem’s activity. By topping up the concentration of sanfetrinem, we were able to show a greater than expected antimicrobial activity when the drug pressure was maintained over time. Time-kill longitudinal data identified a lag-phase upon sanfetrinem treatment prior to bacterial regrowth, which was linked to the post-antibiotic effect of sanfetrinem (Figure 4D), as previously described [40]. Further time-lapse microscopy studies confirm the biphasic killing dynamic of sanfetrinem observed in the time-kill assays; bacterial cytolysis occurred within the first 8 hours of treatment, followed by a slower kill rate, similar to previously observed with faropenem and meropenem [41]. Constant drug exposure leaves a surviving weakened sub-population of tolerant bacteria with a club-like morphology, indicative of cell wall damage (Figure 5), which could facilitate the activity of companion drugs.

The *in vitro* antimicrobial activity of compounds against *Mtb* is typically performed in synthetic 7H9 media supplemented with ADC or OADC containing BSA and glycerol and dextrose as the main carbon sources. This has proven to be non-representative of *in vivo* activity in some cases [42] and previous *in vitro* studies demonstrated that media composition can affect the activity of some β-lactams [19]; we thus explored the impact of different media compositions (Figure 2). The activity of sanfetrinem was affected by the presence of BSA indicating extensive protein binding, which should be taken into consideration when projecting suitable clinical doses. Interestingly, sanfetrinem was more potent in the presence of cholesterol as the sole carbon source, which could be a more clinically relevant *in vitro* model of activity since cholesterol is the main carbon source used by *Mtb* at the site of infection. In MIC determinations against a panel of clinical isolates, sanfetrinem performed favorably compared to several clinically approved β-lactams, including meropenem. The presence of clavulanate improved the overall activity of sanfetrinem against the clinical panel, with a 4-fold increased activity (Figure 3). This reduced shift in the presence of clavulanic acid, compared to the highly hydrolyzed amoxicillin or meropenem, might suggest that the tricyclic carbapenem backbone is inherently more stable to β-lactamase hydrolysis by BlaC. In our studies, this shift was not observed with the laboratory *Mtb* H37Rv susceptible strain (Figure 2C), but it was clearly observed against a rifampicin-resistant strain (Figure S3). That the effect of clavulanate on the activity of sanfetrinem could be better observed in a rifampicin-resistant strains, it is in agreement with previous reports describing a paradoxical hyper susceptibility of drug resistant *Mtb* to β-lactam antibiotics, which is further enhanced in the presence of clavulanate (Figure S2, Table S2) [43].

Any new potential drug developed for TB treatment needs to be included in a multi-drug regimen. As such, understanding how to better combine sanfetrinem could help to optimize its inclusion in future combinatorial treatments. In order to initiate the identification of potential companion drugs from an *in vitro* microbiological perspective, we performed synergy assays against a panel of clinically approved antibiotics, including several currently used in TB therapy. The interaction profile of sanfetrinem differed from that of meropenem (Figure 6A). This was not unexpected since previous studies had shown that interaction profiles are β-lactam dependent [19]. Importantly, sanfetrinem displayed a strong interaction with amoxicillin. Currently, clavulanate is not commercially available and it needs to be added in clinical studies co-formulated with amoxicillin. The inclusion of amoxicillin/clavulanate together with meropenem has proven more effective against MDR/XDR strains than the compounds alone [44], and this could similarly apply to sanfetrinem. In our study, we observed that the strong interaction between sanfetrinem and amoxicillin against the laboratory strain *Mtb* H37Rv (Figure 6B) was somehow masked in the presence of clavulanate (Figure 7), which might be competing with the β-lactamase blocking activity of sanfetrinem [35]. Nevertheless, the activity of the triple combination was slightly improved and greatly enhanced in combination with rifampicin. We also showed that sanfetrinem strongly interacts with rifampicin (Figure 7). The interaction of β-lactams with rifamycins has been described previously [45] and, specifically, amoxicillin/clavulanate showed prevalent interactions with rifampicin and ethambutol against a panel of drug-susceptible and drug-resistant strains [46]. As an additional benefit, gastrointestinal side effects associated with the use of amoxicillin/clavulanate in prolonged therapy can be minimized within the synergistic interaction by reducing the dose. This approach is currently under evaluation in ongoing clinical trials aiming to reduce the treatment duration for Buruli ulcer, a disease caused by Mycobacterium ulcerans (NCT05169554, PACTR202209521256638), by co-administration of amoxicillin/clavulanate to current rifampicin plus clarithromycin WHO recommended therapy [47].

Before embarking on a clinical study in TB, it is expected that a compound demonstrates *in vivo* efficacy in preclinical models. Unfortunately, β-lactams are notoriously difficult to test in rodents due to the high expression levels of DHP-1 enzymes which, like β-lactamases, hydrolyze the β-lactam core before the drug is able to reach the site of action [7]. To quantify the *in vivo* activity of sanfetrinem we employed a DHP-1 knockout model recently developed for the specific purpose of testing β-lactams against TB [24]. Impressively, both sanfetrinem administered subcutaneously and sanfetrinem cilexetil dosed orally demonstrated similar efficacy compared to meropenem or faropenem co-dosed with clavulanic acid (Figure 8). Levels of *in vivo* activity were comparable to other β-lactam studies [29, 30]. Thus, two independent murine experiments confirmed the *in vivo* efficacy of sanfetrinem against M. tuberculosis, similar to other β-lactams already validated in clinical trials.

The concept of drug repurposing can be attractive in any therapeutic area, but particularly in the Global Health space with its limited opportunities for commercial return on investment. The opportunity to circumvent the protracted and expensive pre-clinical and early clinical development phases, proceeding directly to patient efficacy studies, is invaluable. Naturally, hurdles remain in any repurposing approach, particularly when the development program has been dormant for decades. In the case of sanfetrinem cilexetil, which showed variable bioavailability in previous clinical studies (GSK, unpublished data), a significant doubt remains as to whether sufficient lung exposure can be achieved and maintained to have an effect in TB treatment. The *in vitro* potencies against *Mtb* are lower than those observed for many of the Gram-negative and Gram-positive organisms that were targeted in previous Phase 2 studies. However, the *Mtb* potencies are likely underestimated due to the instability of sanfetrinem in longer MIC assays for slow growing bacteria. Additionally, the activity of sanfetrinem could be improved if dosed within a synergistic combination. The data presented in this paper, together with the robust pre-clinical and clinical package from the 1990s, were considered sufficient to justify a proof-of-concept clinical study. A Phase 2a EBA study of sanfetrinem cilexetil is currently underway in South Africa (NCT05388448).

## Acknowledgements

We would like to acknowledge analytical support by Pablo Gamallo from GSK and Carlos Martín, from the University of Zaragoza, for the kind gift of the M. tuberculosis H37Rv and M. bovis BCG derived strains. We are also grateful to the many current and former GSK employees who have supported the clinical re-purposing effort.

## Author contribution

CRediT (Contributor Roles Taxonomy) has been applied for author contribution. Conceptualization: SRG, RGdR, AML, SFB and RHB; Data curation: SRG and RGdR; Formal Analysis: SRG, JR and RGdR; Funding acquisition: SRG; Investigation: SRG, RGdR, MPAC, HB, SA and ASV; Methodology: SRG, RGdR and JR; Project administration: SGR, RGdR, SFB and CJT; Resources: HB, EPdF, MCI and EPH; Supervision: SRG, RHB, SFB, CJT and DBA; Visualization: SRG, RGdR and JR; Writing – original draft: SRG, RGdR and RHB; Writing – review & editing: AML, SFB, CJT and DBA.

## Data availability statement

All data pertaining to this work is within the main manuscript or supplementary information. Primary data are available from the corresponding author upon request.

