## SUPPLEMENTARY INFORMATION for "Sanfetrinem, an oral β-lactam antibiotic repurposed for the treatment of tuberculosis"

^&^ Current address: Certest Biotec, San Mateo de Gállego, Zaragoza, Spain

^$^ Current address: Dep. Bioengineering, University Carlos III of Madrid, Spain

**SUPPLEMENTARY TABLES**

**Table S1. Composition of the GSK in-house library of unique beta-lactams.** The different beta-lactams are classified in sub-families. Hits indicate those compounds active in dose-response studies with a 90% inhibition cut-off at least at 20 µM. Hit discovery rates are indicated globally for the library and for each sub-family of beta-lactams.

**L**

**I**

**B**

**R**

**A**

**R**

**Y**

**C**

**O**

**M**

**P**

**O**

**S**

**I**

**T**

**I**

**O**

**N**

**G**

**S**

**K**

**l**

**i**

**b**

**r**

**a**

**r**

**y**

**H**

**i**

**t**

**s**

**H**

**i**

**t**

**r**

**a**

**t**

**e**

**C**

**e**

**p**

**h**

**e**

**m**

**s**

1

2

9

3

3

7

2

.

8

6

**C**

**e**

**p**

**h**

**e**

**m**

**-**

**l**

**i**

**k**

**e**

7

3

0

0

**C**

**a**

**r**

**b**

**a**

**c**

**e**

**p**

**h**

**e**

**m**

**s**

5

0

0

**O**

**x**

**a**

**c**

**e**

**p**

**h**

**e**

**m**

**s**

5

0

0

**P**

**e**

**n**

**a**

**m**

**s**

1

5

1

4

2

.

6

5

**C**

**a**

**r**

**b**

**a**

**p**

**e**

**n**

**a**

**m**

**s**

4

0

0

**O**

**x**

**a**

**p**

**e**

**n**

**a**

**m**

**s**

3

7

0

0

**P**

**e**

**n**

**e**

**m**

**s**

3

7

1

2

.

7

0

**C**

**a**

**r**

**b**

**a**

**p**

**e**

**n**

**e**

**m**

**s**

8

1

1

2

.

5

**M**

**o**

**n**

**o**

**b**

**a**

**t**

**a**

**m**

**s**

**&**

**o**

**t**

**h**

**e**

**r**

**s**

3

6

0

2

0

.

5

6

**T**

**o**

**t**

**a**

**l**

**:**

**1**

**9**

**7**

**3**

**4**

**5**

**2**

**.**

**2**

**8**

**Table S2.** **Antimicrobial activities of sanfetrinem and other beta-lactams against *M. tuberculosis* strains, including drug resistant clinical isolates.** Information in this table supports Figure 3.

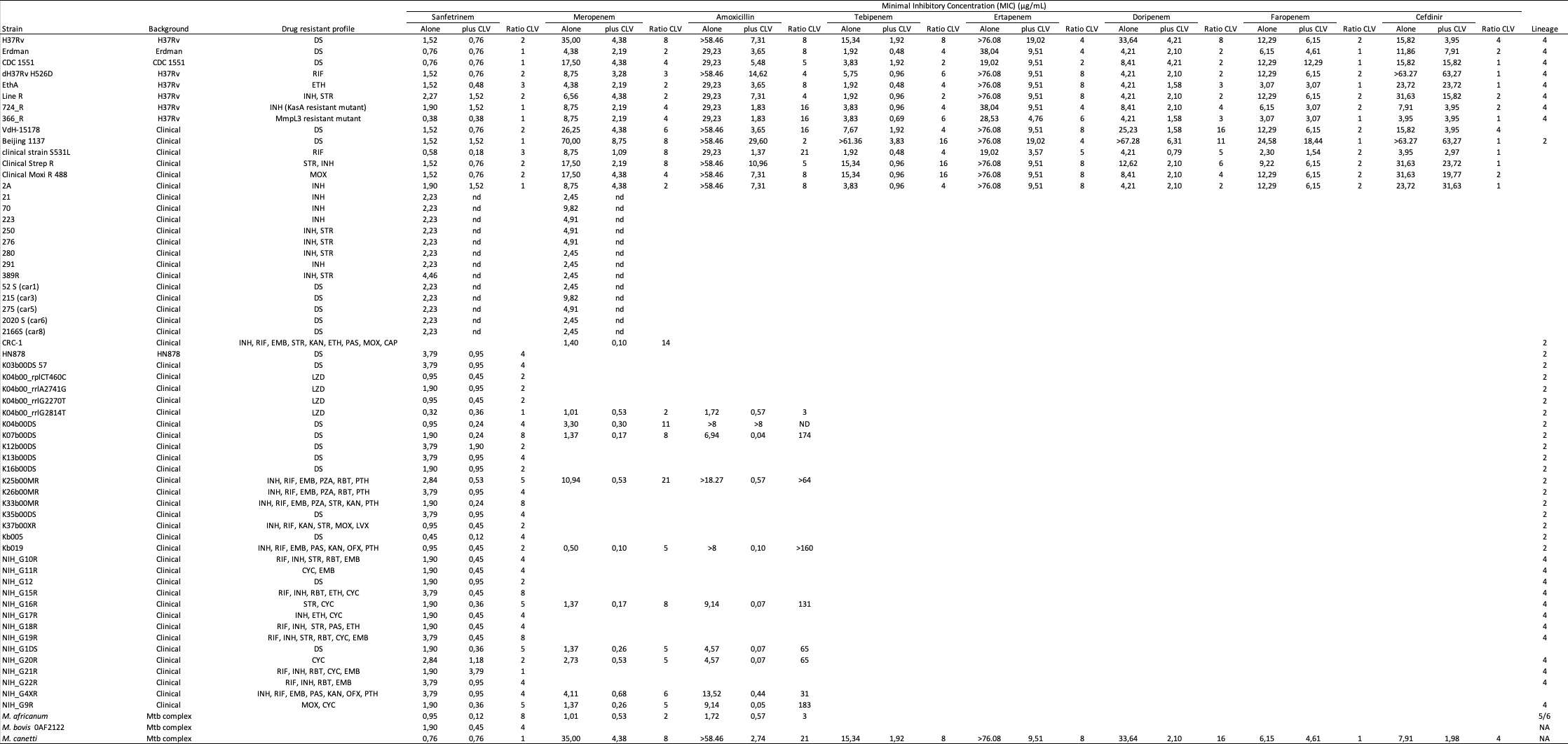

^a^Mycobacterial strains were assayed in liquid 7H9 media containing 0.2% glycerol, 10% ADC supplement and 0.05% Tween80 or tyloxapol. The MIC was determined as the concentration that inhibited growth compared to control wells. See Material & Methods for more detailed technical information. ^b^The concentration of clavulanate used in strains #1 to #14 was 4 µg/mL and in strains #28 to #63 was 25 µM (5.88 µg/mL). DS, drug susceptible; DR: Drug Resistant; MDR: Multidrug resistant; XDR: Extensively drug resistant; CVL, clavulanic acid; CYC, cycloserine; CAP, capreomycin; EMB, ethambutol; ETH, ethionamide; INH, isoniazid; KAN, kanamycin; LVX, levofloxacin; LZD, linezolid; MOX, moxifloxacin; OFX, ofloxacin; PAS, *p*-aminosalicilate; PTH, prothionamide; PZA, pyrazinamide; RBT, rifabutin; RIF, rifampicin; STR, streptomycin.

**SUPPLEMENTARY FIGURES**

**Figure S1. Intracellular dose response assays of sanfetrinem and clinically approved beta-lactams against *M. tuberculosis.*** SFT, sanfetrinem; AMX, amoxicillin; CDN, cefdinir; CFX, cefadroxil; FAR, faropenem; MER, meropenem; BLM, beta-lactam; RLU, relative light units.

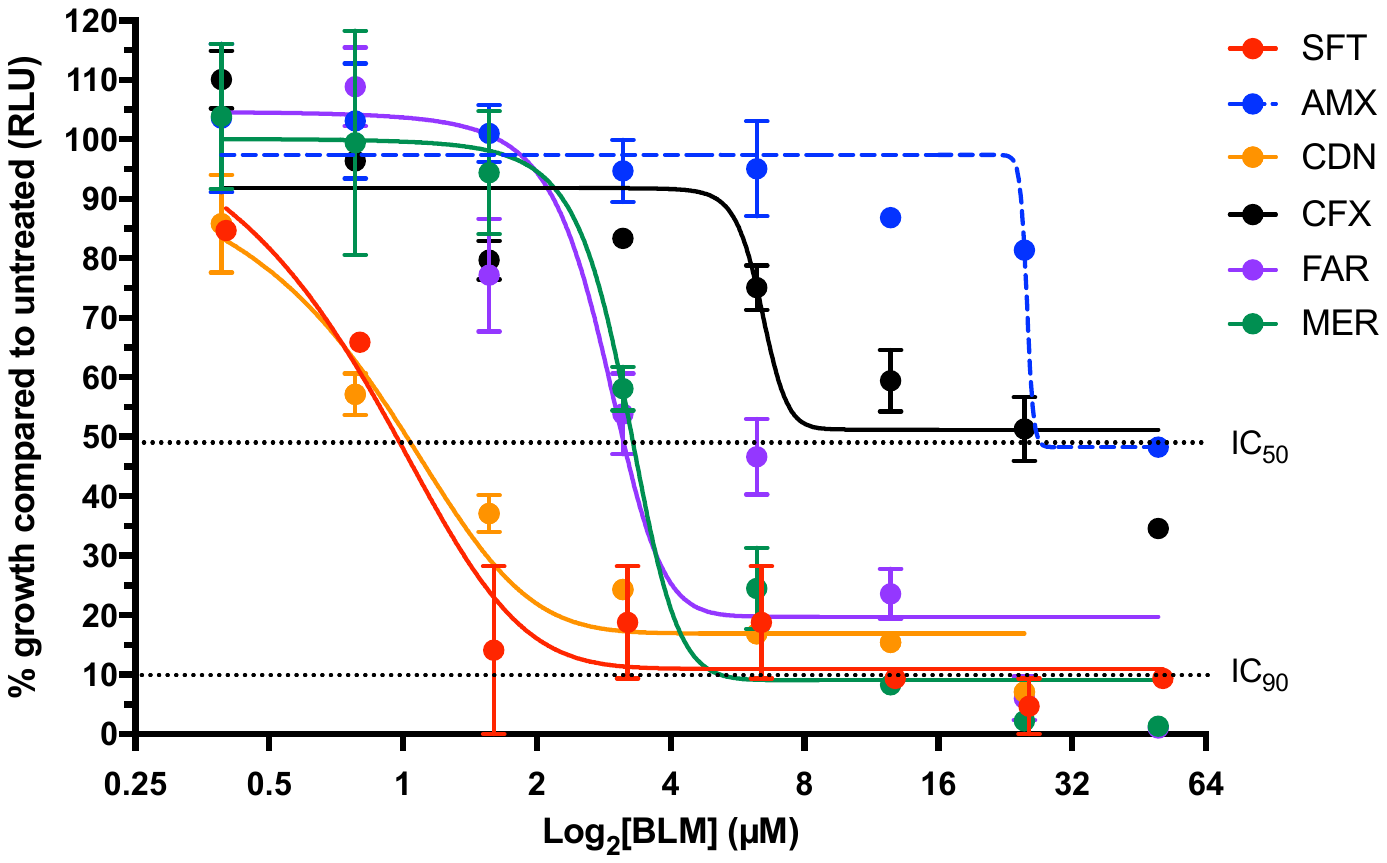

**Figure S2. *In vitro* characterization of different beta-lactams against *M. tuberculosis.*** Dose response studies of amoxicillin, cefadroxil, cefdinir, faropenem and meropenem in standard 7H9 medium (7H9), 7H9 plus the detergent tyloxapol (7H9 + Tx), and 7H9 salts containing cholesterol as the only carbon source (CHO+Tx).

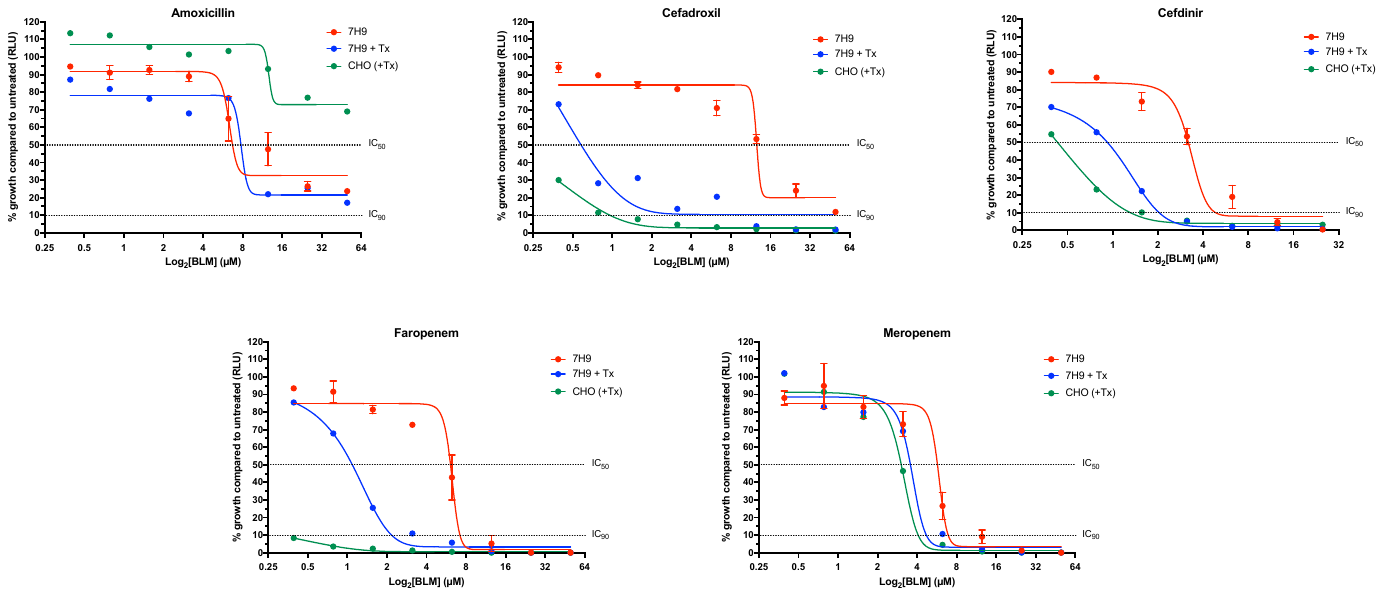

**Figure S3. Time-kill assay of the interaction between sanfetrinem and clavulanic acid against a rifampicin-resistant *M. tuberculosis* strain.** Sanfetrinem (S) and clavulanic acid (V) were tested alone and in combination at 2.5 µg/mL and 5 µg/mL, respectively. At every time point, samples were taken and colony forming units enumerated.

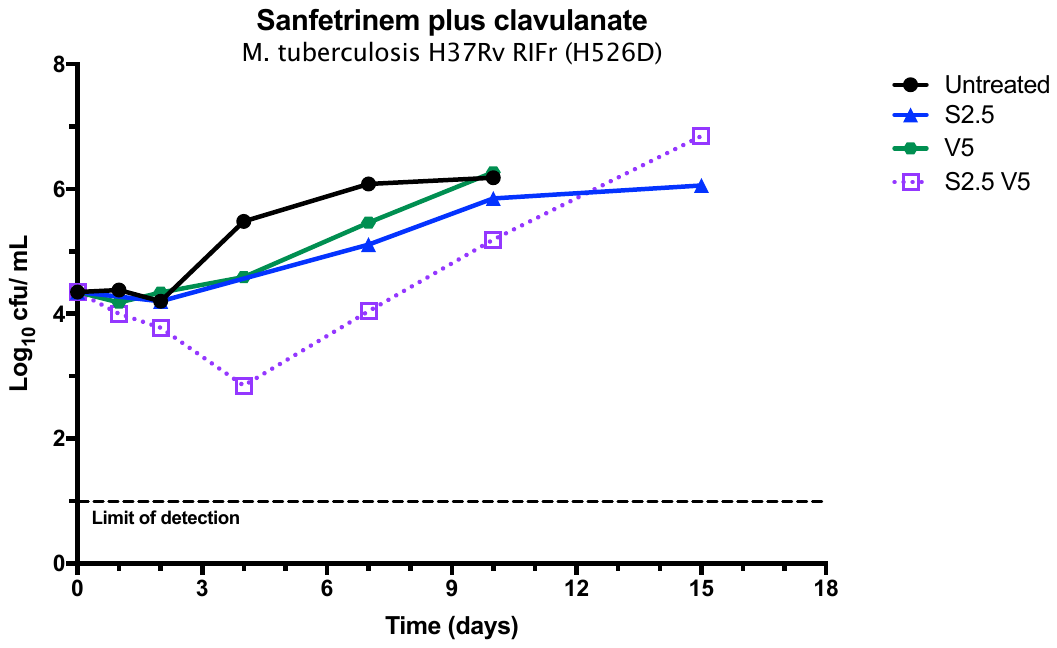

**Figure S4. Post-Antibiotic Effect studies of several beta-lactams against *M. tuberculosis.* (A)** The inoculum effect on the time-to-positivity (TPP) was measured by adding different number of cells (from 10^2^ to 10^5^) to MGIT tubes and growth being continuously monitored. **(B)** Cells treated at the corresponding drug concentrations for 2 hours were transferred (10^5^ cells) to MGIT tubes and the TTP measured. The 2-hours treatment did not affect CFU counts. Rifampicin was included as positive control. Numbers in legend indicate drug concentration in µg/mL for rifampicin, amoxicillin, faropenem, cephradine and clavulanate and in µM for meropenem (5 µM equals to ca. 2.2 µg/mL). TTP, time to positivity to reach a GI > 75. Delta indicates the increase in TTP compared to untreated. AMX, amoxicillin; CLV, clavulanate; CPD, cephradine; FAR, faropenem; MER, meropenem; RIF, rifampicin.

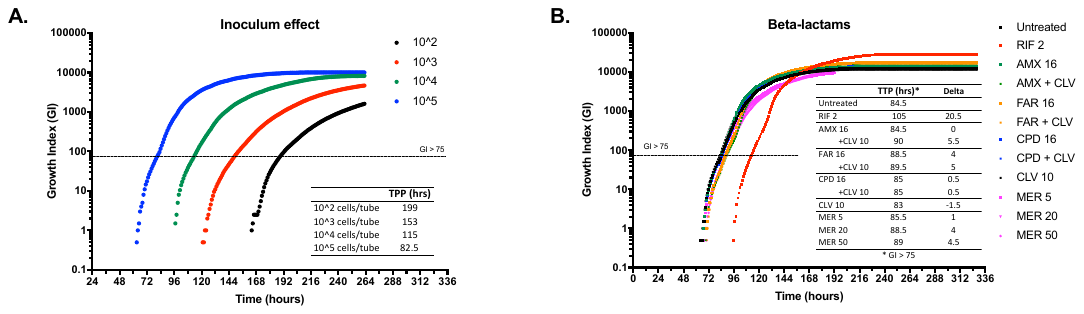

**Figure S5. Pharmacokinetic profiles of sanfetrinem (SNF) and sanfetrinem cilexetil (SNFc) in mice.**

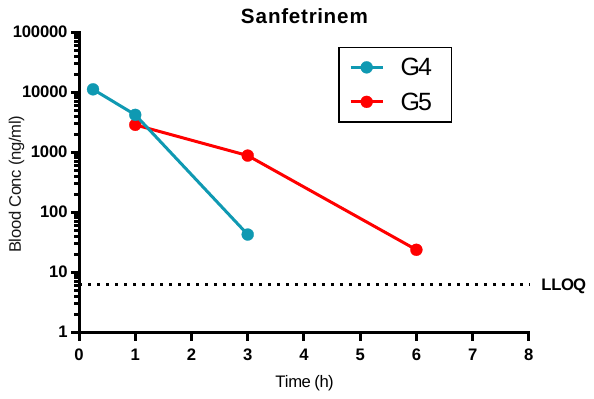

| **Group** | **Compounds**  **Administered** | **Compound**  **Determined** | **T_max_**  **(h)** | **C_max_**  **(ng/mL)** | **C_max_D_**  **(kg*ng/mL/mg)** | **T_last_**  **(h)** | **C_last_**  **(ng/mL)** | **AUC_last_**  **(h*ng/mL)** | **DNAUC_last_**  **(h*ng/ml) per mg/Kg** |
| --- | --- | --- | --- | --- | --- | --- | --- | --- | --- |
| G4 | SNF 200 mg/kg | SNF | 0.25 | 11200 | 56 | 3 | 43 | 11390 | 57 |
| G5 (*) | SNFc 400 mg/kg (**) | SNF | 1 | 2870 | 12 | 6 | 24 | 6545 | 27 |

(*) sample point at t=0.25h was rejected due to administration issues

(**) SNFc 400 mg/kg ≈ SNF 250 mg/kg
